# Single gene initiates evolution of epithelial architecture and function

**DOI:** 10.1101/2021.05.04.442636

**Authors:** Viola Noeske, Emre Caglayan, Steffen Lemke

## Abstract

Epithelial monolayers are a hallmark of the architecture of metazoan tissues: they provide stability, serve as barriers, and fold into organs. Epithelial cells vary in shape, ranging from flat and spread out to tall and slim. Dynamic epithelial shape changes have been explored in the context of tissue folding, where local cytoskeletal modulations cause epithelial bending and folding. Comparatively little is known about how entire tissues are transformed from a short to tall architecture. Here we show that shape regulation in epithelia can be governed by the activity of a single gene. We use a comparative approach in distantly related flies to experimentally decode the developmental program that directs the formation of columnar epithelia in the blastoderm and thus determines the physiological features of the resulting epithelium. We uncover an evolutionary novel, membrane-associated protein that emerged in flies and triggered a new development program, the cuboidal-to-columnar transformation of epithelial tissues. *slow-as-molasses* (*slam*) encodes a Dia/F-actin regulator that exploits an intrinsic morphological plasticity of cells to transform tissues. Our findings demonstrate that a single, newly emerged factor that amplifies its activity in epithelia provides the basis for adaptation and initiates the evolution of novel developmental programs.

## Main Text

Epithelial tissues fulfill crucial functions in the bodies of all animals ^1^. These tissues are thin, continuous monolayers of cells with an apical-to-basal polarity and lateral intercellular adhesions that provide stability, serve as selective chemical barriers, and fold into organs during development ^2^. Epithelia vary in the size and shape of their cells, which range from flat and spread out to tall and slim ^2–4^. Local and dynamic variations in cell shape are a hallmark of tissue folding, and cells are shaped by actin protrusions, actomyosin contractions, and junctional rearrangements to achieve their functions through the folding of epithelial monolayers ^5,6^.

Cell shapes at the level of tissues globally define epithelial architecture and have long been used as classifiers of tissue types ^2–4^. While their functional significance remains unclear, the cell shapes that characterize tissues may be explained in several ways: by tissue-extrinsic forces that determine cell shape through compression or stretching in the plane of the epithelium ^7–9^, by tissue-intrinsic forces and forms of cytoskeletal regulation inherent to the differentiation of complex cell types ^10–12^, or combinations of both.

We investigate these assumptions by comparing differences in the architectures of homologous epithelia of different species. Sharp variations can be observed, for example, in the shapes of cells making up comparable epithelia in the blastoderm, the first epithelial monolayer to form during embryonic development ^3,13^. In some fly species, this tissue is composed of cube-like cells that are about as wide as they are tall, while in others the cells are columnar and several times taller than they are wide ^13^. The distribution of the two types of blastoderm architecture across fly species is not random ^13,14^; it can be traced back to a major evolutionary transition that likely had functional consequences. Flies with cuboidal blastoderm cells, such as midges, gnats, and moth flies, represent the ancestral mode of insect development. A substantial portion of their blastoderm eventually differentiates into extraembryonic epithelia that envelope and protect the embryo from desiccation and infection. In flies with a columnar blastoderm, such as *Drosophila* and their close relatives, extraembryonic epithelia cover a smaller area or no longer form around the embryo at all ^15,16^. The fact that these transitions coincide suggest that a columnar epithelium might have enhanced protective properties compared to a corresponding epithelium composed of cuboidal cells. However, this hypothesis has never been tested.

Here we use a comparative approach in distantly related flies to define the developmental processes that guide columnar epithelia formation in the blastoderm. We provide evidence for the mechanism by which this innovation arose during evolution and test the functional consequences of the altered epithelium for the embryo. Our experiments demonstrate that a novel gene arose in flies; it encodes a Dia/F-actin regulator which exploits the intrinsic morphological plasticity of cells to trigger a cuboidal-to-columnar transformation of tissue architecture. Our findings indicate that a single factor can provide a basis for adaptation and lead to the evolution of novel developmental programs.

### Blastoderm epithelia in flies are made of short or tall cells

To reveal the mechanisms that instruct columnar cell architecture and to address the effects on tissue function resulting from changes in cell shape, we compared two experimentally tractable fly species with distinct types of epithelia in their blastoderm. We used *Drosophila melanogaster* as a fly that features a columnar blastoderm epithelium and the midge *Chironomus riparius* as species that represents the ancestral, cuboidal blastoderm architecture ^17,18^ (**Figure 1A,B**). We quantified both differences in cell height and width, and we also addressed their overall shape (true, tubular columns or tapered at the apical or basal ends) (**Figure 1C, Figure 1 supplement 1**). In epithelial cells, the cytoskeletal structure is primarily maintained through cortical actin closely associated with membranes ^2^. Using actin as a proxy to reveal cell outlines, we found that *Chironomus* blastoderm cells were about half as tall as those of *Drosophila* (12.8 +/- 1.0 µm versus 32.3 +/- 0.9 µm, **Figure 1D**), while the width of the two species’ cells did not differ significantly (4.9 +/- 0.7 µm versus 5.1 +/- 0.7 µm, **Figure 1E**). Neither species exhibited any tapering of cells toward the apical side (**Figure 1F**). On the basal side, however, there were differences: *Chironomus* cells tapered toward the basal substrate, meaning they lost contact with neighboring cells, while those of *Drosophila* were columnar and connected (**Figure 1G**). These results showed that the primary differences between the cuboidal cells of the *Chironomus* blastoderm and the columnar *Drosophila* cells were cell height and the degree of contact between cells at their basal ends.

**Figure 1.**
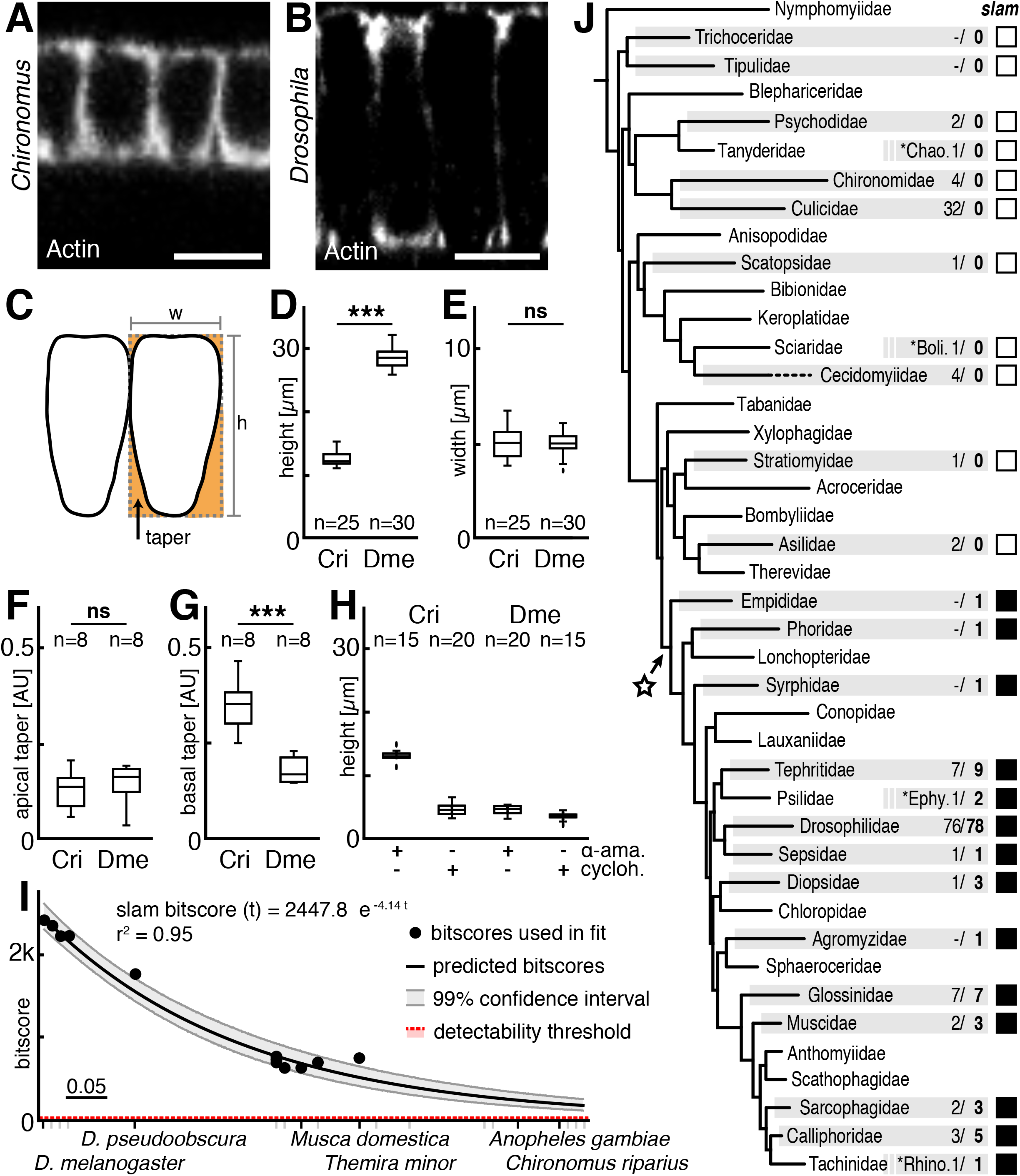
*slam* is a new gene in flies with columnar blastoderm. **A,B**, Blastoderm cells in *Chironomus* (A) and *Drosophila* (B). **C**, Sketch indicating cell width (w), height (h), and taper as measures of blastoderm cell architecture. **D-G**, Comparisons of height (D; *** *P*<0.0001), width (E; ns. *P*=0.76), apical (F; ns. *P*=0.73) and basal taper (G; *** *P*<0.0001) indicate differences in *Chironomus* cuboidal (Cri) and *Drosophila* columnar (Dme) cell architecture. **H**, Blocking transcription (α-ama.) impairs blastoderm formation in *Drosophila*, not *Chironomus*; blocking translation (cycloh.) impairs both. **I**, Extrapolation of sequence divergence in schizophoran *slam* orthologs indicates 100% tblastn detection probability for all flies. **J**, Phylogeny of major fly families, with selected minor families added (asterisk; Chaoboridae, Bolitophilidae, Rhinophoridae) ^28^. Families with sequenced genomes in individual species indicated (grey bar). Indicated counts of high quality genomes (light), genomes with identified *slam* ortholog (bold), character state of *slam* in individual families (open if absent, filled if present), and *slam* emergence (star). P-values calculated with t-test, scale bars 10 μm.

To confirm that both epithelia arise in comparable developmental processes, we investigated blastoderm formation and the first hours of pre-blastoderm development in *Chironomus* (**Figure 1 supplement 2A,B)**. The *Chironomus* embryo initially developed as a syncytium with a total of twelve nuclear divisions of the zygote without cytokinesis. Most nuclei had translocated from the center of the yolk to the periphery of the cell after nuclear division cycle 7. After the last syncytial nuclear division, the blastoderm formed through invaginations of plasma membrane flowing in between peripheral nuclei (**Figure 1 supplement 2C,D**). Formation of the *Chironomus* blastoderm took approximately 120 minutes - about 2-3 times as long as in *Drosophila* (**Figure 1 supplement 2D**) ^19,20^. Additionally, the *Chironomus* blastoderm consisted of fewer cells, which was likely due to the smaller size of its eggs, as the overall cell density in *Chironomus* and *Drosophila* blastoderm epithelia are comparable ^17^. These findings indicate that the differences we observed in cell shape likely arise through genetic regulation of blastoderm development, rather than, e.g., mechanical forces within the plane of the epithelium that could squeeze cells to increase their height at the expense of their width.

### *slam*, a gene known to be required for *Drosophila* blastoderm formation, is present only in flies with tall cells

Columnar blastoderm development in *Drosophila* is genetically regulated by maternal factors introduced as proteins or RNAs, which are translated in the egg, as well as newly transcribed zygotic genes. It was unclear whether this is also true of the cuboidal *Chironomus* blastoderm. To test this, we used specific toxins to inhibit translation or transcription. We found that without translation, neither *Chironomus* nor *Drosophila* developed a blastoderm; without transcription, *Drosophila* blastoderm formation failed but in *Chironomus,* the blastoderm still formed (**Figure 1H, Figure 1 supplement 2E**). These results indicate fundamental differences in the developmental processes that establish blastoderm epithelia in *Drosophila* and *Chironomus*, with columnar blastoderm formation in *Drosophila* depending on the activity of at least one zygotic transcript that is not required for cuboidal blastoderm formation in *Chironomus*.

To identify genes that might encode such a zygotic transcript, we made use of an existing, carefully staged microarray dataset that specifically sampled developmental stages before, during, and after blastoderm formation in *Drosophila* ^21^. Based on this dataset, we compiled a list of candidates that were significantly upregulated during *Drosophila* blastoderm formation (**Figure 1 supplement 3**). We then analyzed a *Chironomus* RNAseq dataset that covered all stages of blastoderm formation, and all candidates for which possible orthologs might be expressed during *Chironomus* blastoderm formation were removed (**Figure 1 supplement 3**). This left a total of 26 genes transcribed during blastoderm formation in *Drosophila* but not in *Chironomus*, including three out of four zygotic genes previously identified as being required for blastoderm formation in *Drosophila* ^22–27^.

Of those 26 candidates, only one appeared to have a *Chironomus* ortholog. This could indicate that the remaining 25 candidates are lineage restricted genes that arose after the divergence of the two species, one of which might explain the differences in *Drosophila* and *Chironomus* blastoderm formation. Alternative hypotheses also existed: we might have been unable to identify orthologs simply because of a poor genome assembly for *Chironomus* or because it had lost the orthologs over time. To adjudicate between the three hypotheses, we performed assembly quality checks and interrogated all 171 of the currently available fly genomes for the presence of our candidates. BUSCO-based testing of genome completeness substantiated a high quality for the assembly of *Chironomus* and the vast majority of other flies (**Figure 1 supplement 4A**), which makes it unlikely that genes would be missed due to poor assembly. Additionally, we could not find orthologs for our candidate genes in other species that represented basal branches of the fly phylogeny. This made it highly unlikely that the genes had been lost specifically in the *Chironomus* lineage.

Instead, our results revealed a quite unexpected pattern in the distribution of specific genes among the species: many of our candidates appeared consistently in the genomes of well-defined subgroups of flies but were entirely absent in outgroups. This suggests that the genes originally arose in flies and that their mapping onto the phylogeny could be used to deduce the stem groups in which they originated (**Figure 1 supplement 4B**). This phylogenetic signal ran deepest for a gene known as *slow-as-molasses* (*slam*) in *Drosophila*. Orthologs of *slam* could be found in all high-quality genome assemblies of Cyclorrhapha and Empidoidea but in none of the other species. This singled out *slam* as very likely the first of these novel genes to have originated in flies, in the stem group of Eremoneura (**Figure 1J**, **Figure 1 supplement 4B**).

When we attempted to identify the ancestral version of *slam*, we were unable to find a sequence with significant similarity. This fact and the way that *slam* maps onto the phylogeny suggest that the gene appeared presumably *de novo* during dipteran evolution. An alternative explanation might be that *slam* had been evolving at a rate that made it hard to detect in non-eremoneuran flies. To clarify this, we sampled species of a group of flies known to have undergone particularly fast sequence evolution, Schizophora ^28^, and extrapolated the probability of detecting putative *slam* orthologs in blast searches of other fly genomes ^29^. These analyses consistently suggested that if *slam* had been present in some ancestral form, the probability of its detection would be 100%, and it should be detectable even in the most basal branching fly families (**Figure 1I**). This meant that our failure to detect *slam* orthologs was not simply due to a very rapid evolution of their sequences.

Another striking finding was that *slam* had been conserved in every one of the fully sequenced fly genomes, which was not the case for any of the other 25 candidate genes (**Figure 1 supplement 5**). This suggested that *slam* has a critical function. We concluded that *slam* was the first of a set of new genes that emerged in eremoneuran flies. Its association with tall cell blastoderm architecture and its expression during blastoderm formation in *Drosophila* ^26,27^ made it the best candidate for a gene responsible for the differences in fly blastoderm development (**Figure 1J**).

### Tall cells in *Drosophila* blastoderm form with a basolateral actin pool, short cells in *Chironomus* without

Next, we attempted to identify mechanisms that might functionally connect *slam*-based blastoderm development to a columnar cell architecture. In *Drosophila*, *slam* encodes a protein of about 135 kDa that is intrinsically disordered, lacking well-defined structural motifs ^26,27^. In the blastoderm, the protein accumulates at the basolateral cell cortex and has been implicated in the regulation of F-actin and the redistribution of plasma membrane ^26,30–32^. Previously, F-actin was shown to play a role in *Drosophila* blastoderm formation ^33,34^. In *Drosophila* embryos that lack *slam* activity, cortical F-actin fails to reorganize into a basolateral scaffold and the blastoderm epithelium does not form ^35^. This indicates that *slam*-based formation of a columnar *Drosophila* blastoderm employs specific mechanisms to modulate local actin dynamics.

To assess the role of F-actin in forming a cuboidal blastoderm in *Chironomus*, we applied two highly specific toxins known to affect the F-actin network in a wide range of animals: Cytochalasin D to disrupt, and phallacidin to stabilize F-actin. Injecting either toxin into *Chironomus* embryos impaired the formation of the blastoderm epithelium (**Figure 1 supplement 6A**). These results confirmed that dynamic F-actin organization is also important for *Chironomus* blastoderm formation. To test whether levels of F-actin influence cell morphology at this stage, we injected actin monomers into *Chironomus* embryos prior to blastoderm formation. Increased F-actin levels did not alter cell height compared to water-injected controls (12.5 +/- 2.0 µm versus 12.8 +/-1.8 µm; **Figure 1 supplement 6B,C**) indicating that actin levels are not responsible for differences in blastoderm formation.

We next asked whether blastoderm architecture was correlated with specific differences in the distribution of F-actin by comparing its localization in the cuboidal blastoderm of *Chironomus* and the columnar blastoderm of *Drosophila*. We used Lifeact-mCherry as a reporter and monitored the temporal changes in the localization of F-actin (**Figure 2A-D, Figure 2 supplement 1**). In *Chironomus*, F-actin remained in a cortical pool that expanded towards the lateral cell membranes as the blastoderm formed (**Figure 2A,B**). The columnar formation in *Drosophila*, on the other hand, was characterized by notable reorganization of F-actin: about half-way through cellularization, a second, separate pool of F-actin formed in the basal region (**Figure 2C,D**). These results support our model that in a fly embryo expressing *slam*, F-actin undergoes a spatial reorganization that plays a role in the formation of columnar blastoderm epithelia.

**Figure 2.**
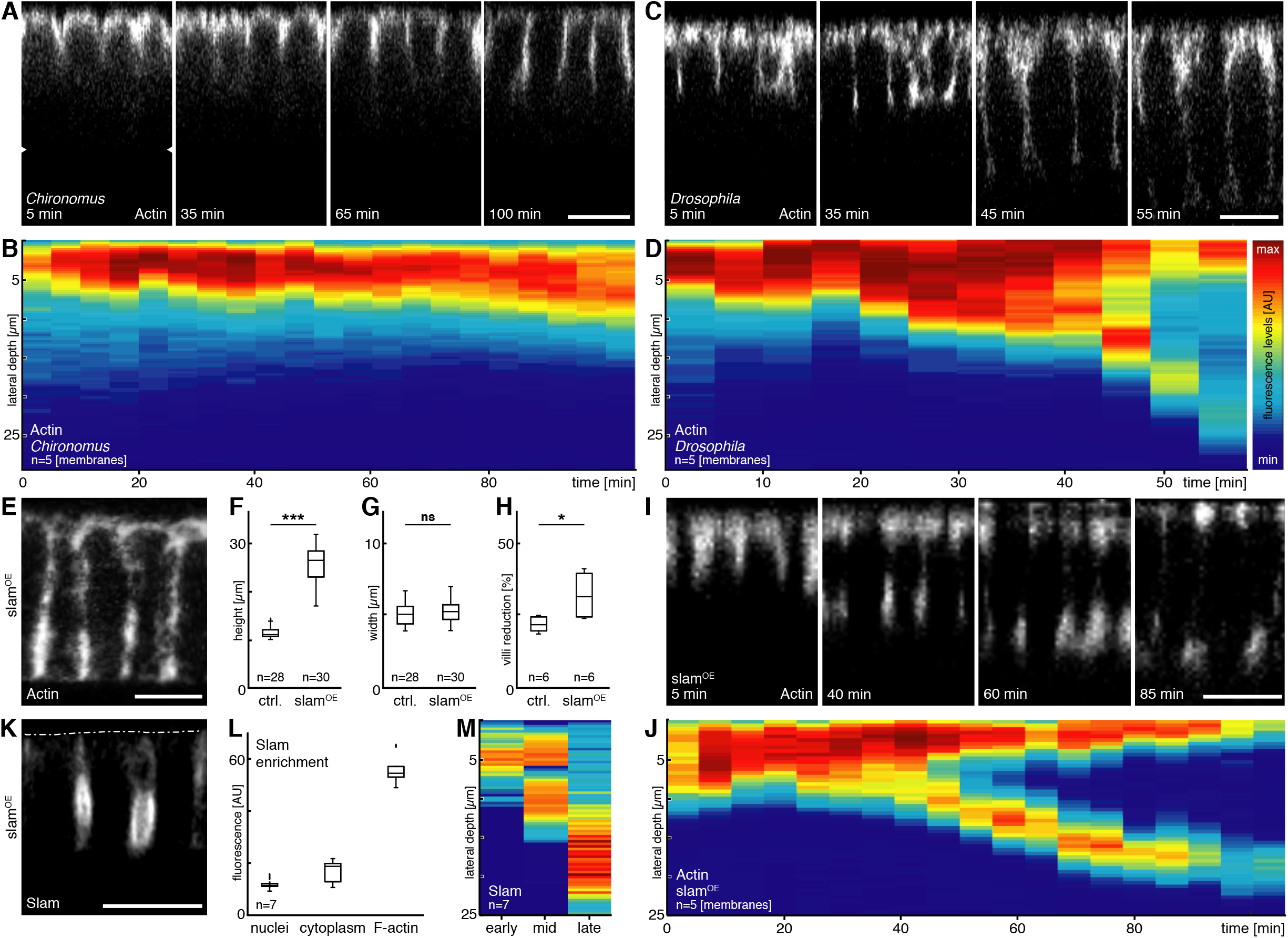
*slam* transforms *Chironomus* blastoderm from cuboidal to columnar. **A-D**, Blastoderm formation in *Chironomus* (A,B) and *Drosophila* (C,D) visualized by the F-actin reporter Lifeact-mCherry. Heatmap kymographs illustrate basolateral distribution of F-actin during blastoderm formation in *Chironomus* (B) and *Drosophila* (D). **E-G**, Expression of *slam* in *Chironomus* embryos (*slam*^OE^) results in tall cells (E) with increased height (F; *** *P*<0.0001) and unchanged width (G; *** *P*<0.0001). **H**, The pool of apical membrane visualized by Gap43-GFP is reduced in *slam*^OE^ embryos. **I,J**, F-actin in *slam*^OE^ embryos with linear cell extension (I) and basal actin (J). **K-M**, Slam-eGFP (K) colocalizes with F-actin (L) and is enriched at basolateral cell faces (M). Indicated are cell apex (dotted line) and imaging depth (triangle). P-values calculated with t-test, scale bars 10 μm.

### Expression of *slam* in *Chironomus* phenocopies *Drosophila* blastoderm architecture

These results raised the question of what would happen to the cuboidal blastoderm formation observed in *Chironomus* if *slam* were present in the midge. To address this, we injected *slam* mRNA into early *Chironomus* embryos (*slam*^OE^ embryos). Strikingly, the blastoderm in *slam*^OE^ embryos now attained a cell height of 28.2 +/- 4.7 µm (**Figure 2E,F**) – more than double the height of wildtype and nearly as tall as cells in the columnar blastoderm of *Drosophila* (**Figure 1B,D**). Notably, this did not increase the width of the cells, which remained unchanged at 5.1 +/- 0.8 µm; it only affected cell height (**Figure 2G**).

For a cell to increase so dramatically its height would require larger amounts of cell membrane. To identify the source, we monitored the supply and demand of cortical membrane in wildtype and *slam*^OE^ embryos by estimating the density of cortical microvilli, as previously established in *Drosophila* ^32,36^. We observed a depletion of the cortical membrane pool in *slam*^OE^ embryos compared to the wildtype pool during blastoderm formation (**Figure 2H**). These results indicate that, potentially, the wildtype has a surplus of cortical membranes that remained unused – but could be redistributed by *slam* activity. In summary, our results indicate that the expression of *slam* is sufficient to initiate a *Drosophila*-like, columnar blastoderm architecture in *Chironomus*.

Next, we wondered whether *Chironomus slam*^OE^ embryos experienced F-actin reorganization comparable to that which occurred in *Drosophila* during blastoderm formation. An examination of the actin cytoskeleton of the embryos revealed that the columnarization of the *Chironomus* blastoderm was accompanied by the appearance of a basal F-actin pool and the establishment of two, seemingly non-continuous apical and basal domains of F-actin resembling what we had observed in *Drosophila* (**Figure 2I,J**). To establish whether this change of F-actin dynamics was associated with a basolateral enrichment of *slam* protein in *Chironomus*, we injected *slam* mRNA encoding an eGFP-fusion reporter. We found eGFP-Slam to be enriched at lateral and basal pools of F-actin (**Figure 2K-M**). This enrichment of the reporter at membranes was similar to the localization of *slam* protein in *Drosophila* ^26^, supporting the conclusion that when *slam* is injected into *Chironomus* embryos, it assumes the functions that it has in *Drosophila*.

Next, we attempted to identify a mechanism linking Slam to basolateral F-actin enrichment in *slam*^OE^ embryos. In *Drosophila*, the basolateral enrichment of F-actin requires the local enrichment of the formin Diaphanous (Dia), a widely conserved F-actin nucleator ^34,37^. This process also requires Rho1, which is activated at basolateral membranes by RhoGEF2 through a direct interaction with Slam ^31^. To indicate sites of F-actin polymerization in *Chironomus slam*^OE^ embryos, we turned to Dia as a reporter. We injected mRNA encoding a previously described GFP-Dia-N reporter to visualize subcellular Dia localization ^38^. In embryos injected solely with mRNA encoding GFP-Dia-N, we observed fluorescence in a single domain located just below the apex of the cuboidal blastoderm (**Figure 3A-C**). In *slam*^OE^ embryos, GFP-Dia-N was observed in two separate pools: one just below the apex and the other at lateral and basal membranes (**Figure 3D,E**). To discern whether this basolateral Dia activity was needed for columnar cell formation, it was necessary to eliminate its activity in a temporally well-defined window, to avoid interfering with any earlier functionality of Dia. We achieved this using a small molecule inhibitor of formin activity (SMIFH2) ^39^, which allowed us to inhibit Dia activity at the onset of blastoderm formation or shortly afterwards. This inhibition of Dia activity reverted the phenotype; it reduced cell height to nearly the wildtype size in SMIFH2-treated *slam*^OE^ embryos. This was consistent with Dia activity during basolateral F-actin polymerization (**Figure 3F**). Taken together, these results suggest that the expression of *slam* increases the localization of Dia to the lateral membrane, which in turn triggers F-actin polymerization at a locally restricted, basolateral site.

**Figure 3.**
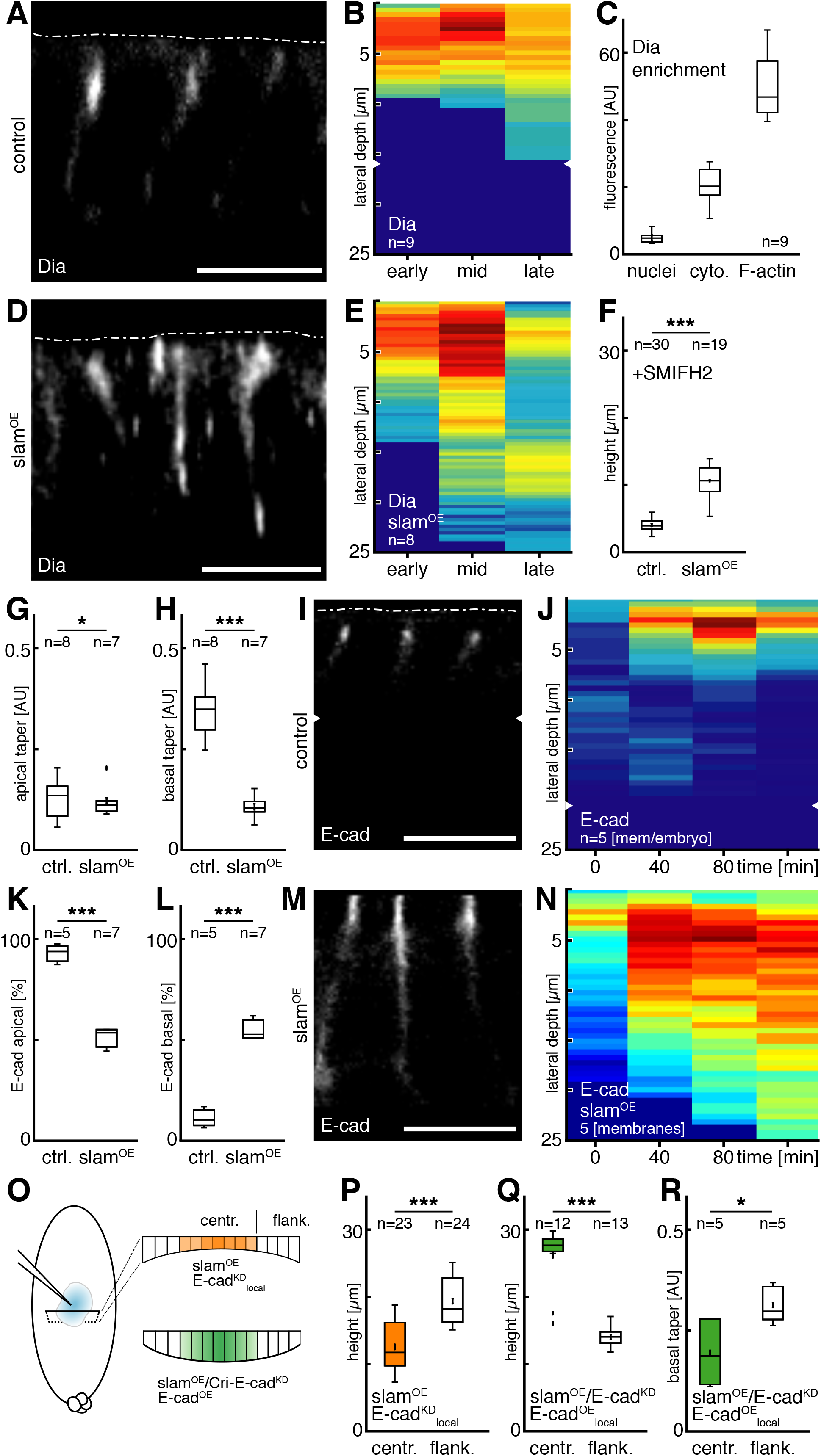
Cell columnarization depends on Diaphanous and E-cadherin for lateral adhesion. **A-C**, Dia-eGFP (A) enriched in a narrow subapical band (B) in colocalization with F-actin (C). **D-E**, In *slam*^OE^ embryos, Dia localizes to subapical and basolateral domains (D) and two separate pools over time (E). **F**, Inhibition of *Dia* activity by injection of SMIFH2 attenuates cell height, which is partially rescued in *slam*^OE^ embryos (\*\*\**P*<0.0001). **G,H**, Lateral cell-cell contact increased in *slam*^OE^ embryos; apical (G; * *P*=0.0401) and basal taper (H; *** *P*<0.0001) is reduced. **I,J**, E-cad-eGFP (I) enriched in narrow subapical domain throughout blastoderm formation (J). **K-N**, Relative E-cad in *slam*^OE^ embryos reduced apical (K) and increase basolateral (L) as reporter (M) distributes over the full basolateral range (N). **O**, Sketch outlining strategy to interrogate *E-cad* function in *slam*^OE^ embryos. **P**, Cell columnarization impaired in *slam*^OE^ embryos in domain of local *E-cad* knockdown (*** *P*<0.0001). **Q,R**, Rescued cell columnarization after global *Chironomus E-cad* knockdown / *slam*^OE^ by local *Drosophila E-cad* activity (Q; \*\*\**P*<0.0001); associated with tighter basal adhesion (R; \**P*=0.024). Indicated are cell apex (dotted line) and imaging depth (triangle). P-values calculated with t-test, scale bars 10 μm.

### Slam extends lateral adhesion and E-cadherin in *Chironomus* blastoderm

While the basolateral accumulation of Dia and F-actin polymerization indicate that *slam* has comparable functions in *Drosophila* and *slam*^OE^ *Chironomus* embryos, the mechanistic links between this local activity and the global elongation of basolateral cells remained unclear. A possible lead came from planar cell-cell dynamics in actively reshaping epithelia, where a model has been proposed in which excess Dia activity generates small patches of stable cortical actin that might immobilize and induce clustering of E-cadherin (E-cad) ^40,41^, a key molecule in cell-cell adhesion in epithelial tissues. In the context of blastoderm formation, F-actin/E-cad patches extending along the apical-to-basal axis of adjacent lateral cell membranes might act as a mechanism effectively “zippering” cells together. By analogy to planar adhesion zippers in cell culture ^42^, this mechanism predicts that lateral cell-cell contact zones selectively expand and could, in an epithelial context, produce taller cells.

To test this prediction, we first investigated whether *slam* activity leads to increased lateral cell-cell contacts by comparing cell tapering in wildtype and *slam*^OE^ embryos. An increase of basolateral cell-cell adhesion should reduce cell tapering, and in fact we found significantly less tapering in *slam*^OE^ embryos than wildtype (**Figure 3G,H**). To test whether the differences in lateral cell-cell adhesion between wildtype and *slam*^OE^ embryos correlated with differences in the distribution of adhesion molecules, we used a *Drosophila* E-cad-GFP reporter to reveal and quantify adherens junctions (AJs). In embryos injected only with mRNA encoding the reporter, AJs were found almost exclusively in the subapical domain (0-5 µm below the apex displays 88.7 +/- 3.5% of all GFP signal, **Figure 3I,J**). In *slam*^OE^ embryos, AJs were still enriched subapically (49.6 +/- 3.7%), but also widely distributed along the basolateral cell membrane (50.4 +/- 3.6%, **Figure 3K-N**).

These results are consistent with the presence of a lateral zipper and a continuous generation of lateral F-actin/E-cad patches as drivers of cell columnarization. Alternatively, Slam might instruct cell shape changes through a different mechanism, with accumulations of lateral E-cad playing only a secondary role. To distinguish between these two models, we assessed the role of basolateral E-cad. E-cad is required during earlier stages of development (**Figure 3 supplement 1A**), so we needed a way to interfere with E-cad specifically during blastoderm formation. For this, we first used a knockdown of *Chironomus E-cadherin* (*Cri*-*E-cad* RNAi_local_), which takes effect late and thus, in a locally restricted manner. This established a central domain of blastoderm formation that started but stalled and a peripheral domain in which complete cells formed (**Figure 3 supplement 1B-F**). We then added *slam* mRNA by injection into the center of the embryo. This yielded *Cri-E-cad* RNAi_local_+*slam*^OE^ embryos with a cell height of 12.6 +/- 3.6 µm in the central domain of the blastoderm, compared to 19.3 +/- 3.4 µm in the peripheral domain (**Figure 3O,P**). Conversely, we analyzed *slam*^OE^ embryos in which *E-cad* was knocked down early and globally (*Cri*-*E-cad* RNAi_global_), and then locally rescued by the injection of mRNA encoding *Drosophila* E-cad-GFP. This rescue in *Cri-E-cad* RNAi_global_ embryos resulted in small islands of cells which could be identified by junctional GFP signals (**Figure 3 supplement 2A-E**). Even in cases where these islands were clearly detached from the peripheral blastoderm cortex and basement, cells would still initiate cell columnarization (**Figure 3O,Q,R**). Taken together, these results indicate that E-cad plays a critical role in blastoderm columnarization and strongly suggest that *slam* functions by basally extending lateral cell-cell adhesion through newly assembled F-actin/E-cad patches.

### A columnar blastoderm reduces the water permeability of embryos

With the completion of blastoderm formation, the fly embryo creates a first epithelial barrier between yolk and environment. In *Chironomus*, we tested the effectiveness of this barrier with respect to water flow by placing embryos into a high salt solution (**Figure 4**). In this setting, the shrinkage of the embryo provides a reasonable proxy for water efflux. Unfertilized eggs do not form a blastoderm; when placed into the salt solution alongside embryos with a blastoderm, the unfertilized eggs shrank substantially faster than the embryos (**Figure 4A,B,D**). This demonstrated that the epithelium attenuates water exchange between yolk and the environment. To address whether a columnar blastoderm reduced water efflux past the epithelial barrier in *Chironomus* embryos, we compared the rates of shrinkage of control and *slam*^OE^ embryos. Strikingly, we found that shrinkage was substantially reduced in *slam*^OE^ embryos, at rates that correlated with the concentration of salt (**Figure 4C,D**). These results indicate that embryos with a columnar blastoderm experience less water loss and provide evidence that a columnar cytoarchitecture increases the barrier function of their epithelia.

**Figure 4.**
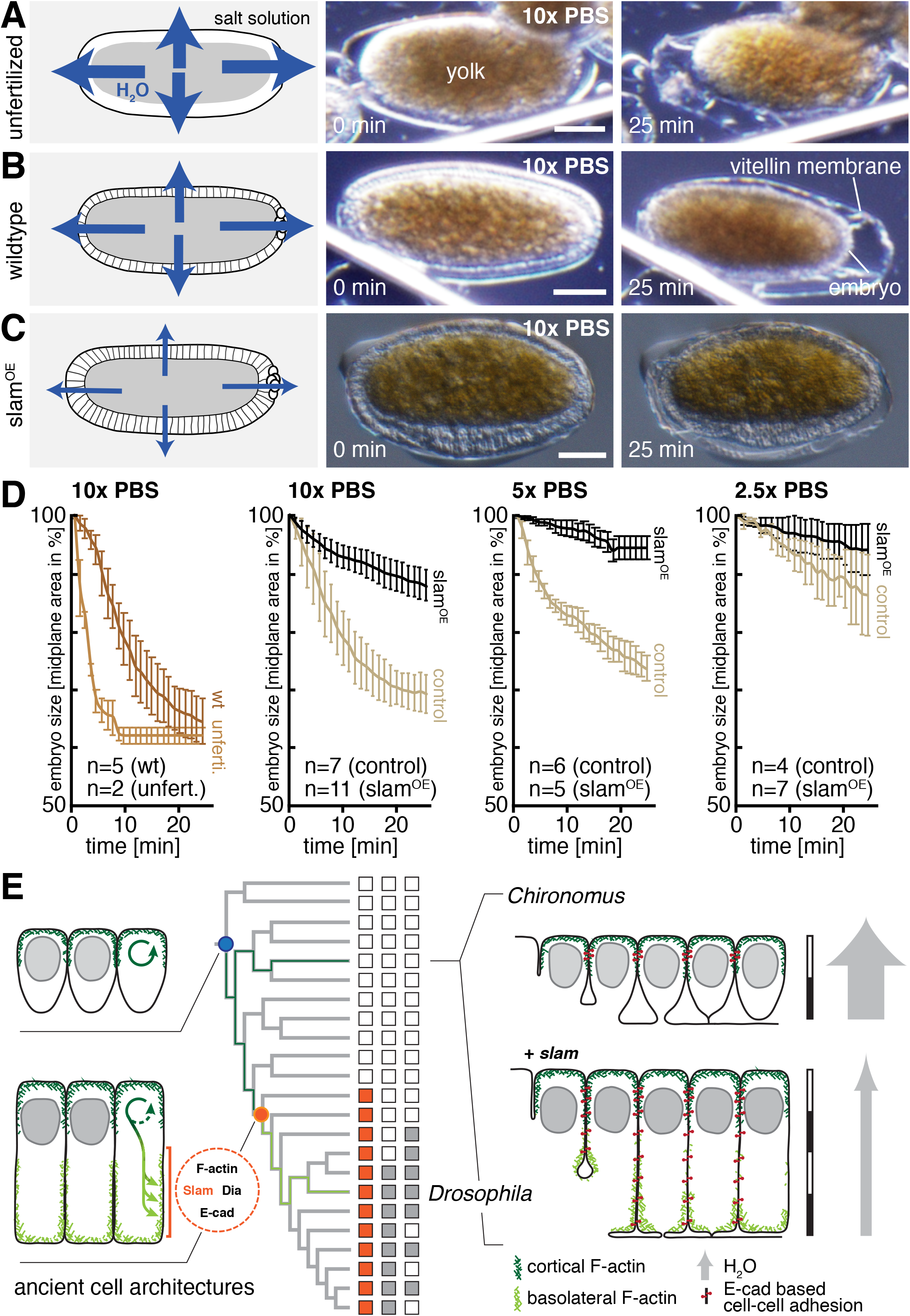
A columnar epithelium constitutes an improved osmotic barrier. **A-C**, Embryo deformation as response to inferred water loss in unfertilized egg cell without blastoderm (A), with a cuboidal (B) and a *slam*-induced columnar blastoderm (C). **D**, Embryo shrinkage is a function of salt concentration and cell height. **E**, Model outlining the evolution of columnar blastoderm formation by *slam* as ‘founder gene’ of a new developmental program. Scale bars 50 μm.

## Discussion

### A model to explain the ancient switch from a cuboidal to a columnar blastoderm

Here, we reveal an astonishingly simple mechanism that is capable of transforming a cuboidal blastoderm into one composed of columnar cells. We demonstrate that this evolutionary transformation very likely originated with the emergence of a single gene, *slam*, and provide evidence that a columnar epithelium represents an enhanced protective barrier that reduces the water permeability of an embryo (**Figure 4E**). Mechanistically, *slam* integrates seamlessly into pre-existing cellular infrastructures; it converts cuboidal cells into columnar ones by drawing on surplus membrane, promoting localized F-actin polymerization and accumulation of E-cadherin to generate patches of lateral cell adhesion. Cells likely elongate in a process analogous to the lateral cell zippering that has been observed in tissue culture ^43^, where patches of lateral adhesion seed a self-organized zipper that tethers and progressively extends zones of lateral contact along apical-to-basal cell interfaces. Alternatively, *slam* might recruit specific factors that promote membrane extension to the basal side of the cell, for example, through guided vesicle fusion ^44^. Irrespective of the exact mode of cell elongation, we show that a local and laterally restricted activation of commonly available cytoskeletal regulators by *slam* creates tall cells, which form a yolk-enclosing columnar epithelium and provide enhanced protection against desiccation.

### Exploitation of existing cytoskeletal regulators to alter cellular morphology

We propose that the emergence of *slam* exploited inherent capacities of cells to change their morphology by remodeling the actin cytoskeleton. The cytoskeleton relies on simple core components that are deeply conserved; its dynamics are controlled in space and time by Rho GTPases, which are regulated by GTPase-activating proteins (GAPs) and guanine exchange factors (GEFs) to act as molecular switches ^45^. This control over actin dynamics is modulated in various ways to result in new cell shapes. In evolutionary terms, Rho GTPases and their regulators have undergone duplications and acquired new functions, representing a potential toolbox to diversify cell morphology. Consistent with this scenario, the metazoan radiation of Rho regulators correlates with a vast divergence in cell shapes and functions ^11,45^. Other mechanisms can have similar effects on actin dynamics and cell shape; they may be modulated by proteins that restrict or localize Rho activity to distinct new subcellular domains ^46,47^. In *Drosophila*, Slam has been proposed to serve as such a guide that localizes with other factors to the basolateral blastoderm membrane, possibly catalyzed by a phase transition with its own mRNA and an intrinsically disordered region in the protein ^30^. At this location, Slam enriches RhoGEF2 and Patj ^31^, a basal scaffolding protein that physically interacts with actin-binding proteins including Moesin, Par-3, Par-6, and Myosin ^48–50^. Regardless of the precise mechanisms by which Slam is enriched at the basolateral site of the cell, its ability to repurpose existing Rho activity and locally modulate the actin cytoskeleton gives this single factor the capacity to induce major morphological changes in inherently plastic cells.

### A “founding gene” drives the emergence of a novel developmental program

New genes have repeatedly arisen over the course of evolution and have contributed to the divergence of specific taxa ^51^. Slam likely emerged approximately 190 MYA. Due to its ability to transform an ancient developmental process, we propose that it acted as the ‘founding gene’ of a novel program. The functionality of this program was enhanced with the emergence of additional genes such as *bottleneck* and *nullo*, which are required for blastoderm formation in *Drosophila* ^23,24^, but which were lost again in specific lineages (**Figure 1 supplement 4B**). In contrast to these presumably supportive factors, *slam* was never lost and has evolved to be the most crucial element for proper blastoderm formation in *Drosophila*; without it, embryos fail to initiate the process ^35^. This suggests that over time, not only did *slam* become the dominant component of blastoderm-formation mechanisms, but ultimately, it replaced the ancient process of cuboidal blastoderm formation in entirety.

The apparent enrichment of new genes in the developmental program guiding columnar blastoderm formation may reflect a common principle to generate novelty. For example, the early embryos of fly and fish have been shown to be enriched for the activity of evolutionarily young genes, many of which have short sequences and can be transcribed in between rapid cycles of nuclear divisions ^52^. We speculate that ‘founding genes’ emerge from this reservoir of new genes and continue to build on it as novel developmental programs are established. This principle differs notably from well-described tinkering in the protein coding sequences of existing genes or in regulatory elements ^53,54^; it demonstrates how a unique developmental program can emerge ‘from scratch’.

### Targeted actin reorganization as a mechanism for the evolution of epithelial architecture

Epithelia fulfill a protective function by limiting the exchange of molecules between the internal body milieu and the external environment. The fly blastoderm represents an interface between yolk and the surroundings of the embryo. When Slam triggers the development of a columnar blastoderm, there is an increase in the osmotic barrier function of this epithelium. We envision that a comparable transformation of the ancient fly blastoderm might be advantageous by reducing the risk of desiccation of early embryos. This could explain why the innovation of *slam* in flies permitted a reduction in the extent of extraembryonic tissues, which are expensive to build in terms of resources but offer protection from desiccation ^16^.

Similar epithelia with columnar cell architecture and boundary function are found in diverse contexts in species and tissues. Examples are organs of the digestive system, both in vertebrates and invertebrates ^55^. Here we show how this type of epithelial structure arose through a relatively simple mechanism in flies, with direct effects on the functions of the tissue. It is tempting to speculate that the columnar cell architecture in tissues at different developmental stages or in other species might have developed through analogous cellular mechanisms, as an interesting example of convergent evolution. Indeed, experiments in cell culture suggest that local remodeling of F-actin could play a general role in cell columnarization ^56^, and the egg chamber in flies or hair follicle cells in vertebrates provide independent examples where local adhesion and spatially restricted activity of molecular switches play a role in maintaining a columnar cell shape ^57,58^. Further investigations are needed to decipher the exact mechanistic basis of columnarization in these cells, and to address the shape-specific function of columnar epithelia as a general factor in the selection of epithelial architecture during animal evolution.

## Conclusion

Our work reveals that embryonic epithelial cells have the capacity to transform their shapes, resulting in novel tissue properties with physiological consequences. We show how this capacity was exploited by a single, new gene, which transformed a cuboidal blastoderm into a columnar one by repurposing existing molecular machinery. Using a fly epithelium as a model, our results provide insight into how newly emerged genes lead to the evolution of a new developmental program and thus adaptation to the environment. More broadly, we propose that novel regulators of the cytoskeletal organization play a key role in the evolution and diversification of tissue architecture.

## Acknowledgements

We thank Maike Fath and Naima Ruhland for assistance with cloning, and Paula González for help with Python. We thank Annika Guse, Lazaro Centanin, Takashi Hiiragi, Marie Jacobovitz, Ingrid Lohmann, Jan Lohmann, Yu-Chiun Wang, Jochen Wittbrodt and members of the Lemke lab for discussion and comments on the manuscript.

## Author Contributions

Performed experiments: VN. Analyzed experiments VN and SL. Sequence analyses EC and SL. Conceived and planned research: VN and SL. Drafted manuscript: VN and SL. All authors read and commented on the manuscript.

## Methods

### Fly keeping

A laboratory culture of *Chironomus riparius* (Meigen, Bergstrom strain) was maintained at 25 °C and a constant 17/7-hour day/night cycle as previously described ^59^. Experiments in *Drosophila melanogaster* were carried out using strain *w1118*.

### Cloning, RNA synthesis and fluorescent protein generation

To generate a template for *in vitro* mRNA synthesis for the full length coding sequence (CDS) of *slam* fused to GFP, a fragment encoding GFP-Slam was amplified by PCR from pMT-GFP-slam ^31^ (gift from J Großhans) using primer pair 5’-GAATACAAGCTTGCTTGTTCTTTTTGCAGAAGCTCAGAATAAACGCTCAACTTTGGCAGATAAAATG/5’-AACAGGTCTCTTCGATCAGACCTCCACGGCCCTCCGGTCCATCAG, digested with *Hind*III and *Sal*I, and ligated in the respective cloning sites in pSP35T as described ^17^. The resulting pSP-GFP-slam was subjected to site-directed mutagenesis with primer pair 5’-CGTGACCACCCTGACCTACGGCGTG/5’-AGGGTGGGCCAGGGCACG to add a GFP^F65L^ mutation that resulted in pSP-eGFP-slam. Capped mRNA of full length *slam* was generated using pCS2-Slam as template ^31^. To visualize membranes, a GAP43-eGFP fusion construct was used. To obtain a template for mRNA synthesis, the Gap43-eGFP coding sequence was generated by in-frame Gibson cloning of the Gap43 encoding sequence, a short linker (GSAGSAAGSGEV), and a previously published pSP vector with 3’-terminal eGFP pSP-Mab-bsg-eGFP ^60^. A fluorescent reporter for subcellular Diaphanous (Dia) localization was generated using the GFP-Dia-N fragment that was described previously ^38^. To obtain the template for mRNA synthesis, a fragment coding for GFP-Dia-N was PCR amplified from GFP-Dia-N-pUAST-attB (gift from B Shilo) using primer pair 5’-AACAGGTCTCACATGGTGAGCAAGGGCGAGGAGCTGTTCAC CGG/5’-AACAGGTCTCTTCGACTACGCCACACCATTAGCCTCCATCAA, digested with *Bsa*I and *Sal*I, and ligated by matching overhangs into *Nco*I/*Sal*I digested pSP35T. The Kozac sequence was optimized using site-directed mutagenesis with primer pair 5’-TTGGC AGATAAAATGGTGAGCAAG/5’-AGTTGAGCGTTTATTCTG, resulting in pSP-GFP-Dia. The template for mRNA encoding a fluorescent reporter for subcellular localization of E-cadherin (E-cad) was generated by in-frame Gibson cloning of fragments encoding E-cad and a short linker (GSAGSAAGSGEV) into a pSP vector carrying a 3’ terminal eGFP CDS. The fragment encoding full length E-cad and linker was amplified from pBabr-5sqh-E-Cad-stf-mRuby3-3sqh (unpublished, gift from YC Wang) using primer pair 5’-CGCTCAACTTTGGCAGATAAAATGTCCACCAGTGTCCAGCGAATGTC/5’-CCTCGCCCTTGCTCACCATCACCTCGCCGGAGCCGGC; the pSP backbone including eGFP was amplified using 5’-CCATGGTGAGCA AGGGCG/5’-TTTATCTGCCAAAGTTGAGC. RNA was *in vitro* transcribed using SP6 Polymerase (Roche). Capping and polyA-tailing was performed using ScriptCAP 1 Capping System and Poly(A) Polymerase Tailing Kit (CellScript). A *Chironomus* ortholog of *E-cad* (*Cri-E-cad*, GenBank_XXXXXX) was identified from transcriptome sequences and cloned after PCR amplification from cDNA. The template to generate *Cri-E-cad* double-stranded RNA (dsRNA) comprised pos. 2677 to 3695 (pos. 1 refers to first nucleotide in ORF). The fragment was amplified by PCR with primer pair 5’-TAATACGACTCACTATAGGGAGACCACGCTGTTGACAAGAGCGGATCGA/5’-TAATACGACTCACTATAGGGAGACCACCCTCGCATTGTTGGCGCATATG with included T7 promotors; dsRNA synthesis was carried out as described ^17^. Lifeact-mCherry was generated as a recombinant protein as described ^60^.

### Immunohistochemistry

Embryos were fixed for 40 minutes in a mixture of 8.2% formaldehyde in PBS (137 mM NaCl, 2.7 mM KCl, 10 mM Na_2_HPO_4_, 2 mM KH_2_PO_4_, pH 7.4) covered with n-heptane. Devitellinization was carried out by shaking embryos in equal of ethanol (95%) and n-heptane, followed by washing (ethanol, 90%) and storage in 90% ethanol at 4 °C. Staining of DNA and F-actin were carried out as described with minor modifications ^17,60^: the phallacidin stock (200 units/ml, Invitrogen B607) was diluted in PBS (1:50), and embryos were stained for three hours at room temperature. DNA was stained using 4’,6-diamidino-2-phenylindole (DAPI, Life Technology D1306) at a final concentration of 5 µg/ml for one hour, followed by several washes in PBS. To reveal the localization of eGFP-fusion proteins in fixed tissues, embryos were injected with mRNA encoding the respective reporter at pole cell stage. Following fixation and methanol-free devitellinization, antibody staining against GFP was carried out essentially as described ^17^. Briefly, embryos were rehydrated, washed, blocked with 5% blocking solution (Roche, 11 921 637 001) in PBT for at least two hours at room temperature, and then incubated with primary antibody (1:250, anti-GFP chicken IgY unconjugated 2mg/ml; Life Technologies, A10262) in 5 % blocking solution overnight at 4 °C. Embryos were washed and incubated with secondary antibody (1:250, donkey anti-chicken Alexa 594, Jackson, 703-505-155) for 3.5 hours at room temperature. Embryos were then washed, transferred to glycerol/PBS and mounted for imaging.

### Injections of small compound inhibitors, mRNA, dsRNA, and recombinant protein

Embryos were collected, prepared for injection, and injected essentially as described ^17^. Small compound inhibitors were used at concentrations as previously described: α-amanitin (50 µg/µl ^61^), cycloheximide (0.5 mM ^61^), SMIFH2 (100 nM ^39^) and Cytochalasin D (2mg/ml). α-amanitin and cycloheximide were injected at onset of blastoderm formation. Phallacidin and Cytochalasin D were injected at syncytial blastoderm stage or at the onset of blastoderm formation as indicated; SMIFH2 was injected at the onset of blastoderm formation. Unless indicated otherwise, injections of capped mRNAs were performed at pole cell stage (about two to three hours after egg lay) with the following concentrations: *slam-eGFP* (2.2 µg/µl) *GAP43-GFP* (1.7 µg/µl) and *Dme-E-cad-GFP* (2.0 µg/µl). Recombinant Lifeact-mCherry (2.0 µg/µl) and TexasRed-Histone H1 (0.7 µg/µl) were generated as described ^60^, and injections were performed before last nuclear division (about six hours after egg lay, Lifeact-mCherry) and at pole cell stage (about two hours after egg lay, TexasRed-H1), respectively. For global knockdown of *E-cad*, *Cri-E-cad* dsRNA was injected at nuclei migration stage, for local knockdown at late nuclei migration stage (after last nuclear division); concentration was 0.8 µg/µl. Restricted *Cri-E-cad* knockdown was established by a time series experiment in analogy to previously published work ^62^. Briefly, *Cri-E-cad* dsRNA was injected into the center of the embryo. If injection was carried out before onset of nuclear migration (corresponding to three hours after deposition and four hours prior to onset of blastoderm formation at 25 °C), cellularization did not start anywhere in the embryo and peripheral nuclear divisions were blocked. Restricted knockdown was then defined such that blastoderm formation continued and completed at the poles, but not the injection area in the center of the embryo. Ideal timing for E-cad dsRNA injection for global KD was thus determined to be at five hours after deposition (corresponding to two hours prior to onset of blastoderm formation at 25 °C). Local rescue with E-cadherin-eGFP in Cri-E-cad RNAi embryos was carried out by injection after the penultimate syncytial nuclear division, about one hour before onset of blastoderm formation in a 3-to-1 mix with recombinant Lifeact-mCherry.

### Microscopy

Dynamics of blastoderm formation were recorded using DIC and confocal microscopy. Embryos were imaged within their central third along their anterior-to-posterior axis without random orientation along the dorsoventral axis. Time lapse recordings with DIC were used to analyze progression of cellularization in embryos injected with α-amanitin, cycloheximide, SMIFH2, Cytochalasin D, or phallacidin compared to water injected control embryos. Recordings were taken using a Nikon Eclipse Ti with a 20x objective (Nikon Plan Apo 20x/0.75 OFN25 DIC N2) for 2.5 hours in two minute intervals at the optical median plane of a sagittal section. Time lapse recordings with confocal microscopy were performed to reveal F-actin dynamics via Lifeact-mCherry in otherwise wildtype embryos and embryos that were additionally injected with mRNA for *slam* and *E-cad-eGFP*. Time lapse recordings were taken by single-photon confocal imaging in a Leica system (SP8) using a 63x immersion objective (HC PL APO 63x/1.30 Glyc CORR CS2). Cellularization was recorded in 0.42 µm sections over a range of about 30 µm, with either single or simultaneous detection of mCherry and eGFP. Field of view was 1024×1024 pixels, with voxel size 0.24 x 0.24 x 0.42 µm and volumes were collected at five minute intervals for 2.5 hours. Early embryonic development (two to seven hours after egg lay) were recorded with simultaneous detection of TexasRed (Histone H1) and brightfield. Volumes were collected at six minute intervals for five hours with a voxel size of 0.24 x 0.24 x 2 µm over a z range of about 50 µm. Immunohistochemically labeled embryos were imaged as confocal z-stacks (8bit) using 63x magnification and identical voxel sizes. Unless noted otherwise, images in figure panels are xz-reslices of z-stacks using an average intensity projection of 2.4 µm.

### Embryo shrinking assay

Embryos were lined up, covered with oil, and injected with *slam* mRNA (*slam*^OE^), H_2_O (control), or left without injection (wt or unfertilized eggs). At the end of blastoderm formation, embryos were washed briefly with heptane to remove the halocarbon oil, immersed in 10x PBS, 5x PBS or 2.5x PBS, and embryo behavior was captured in time lapse recordings (every 20 seconds for 25 minutes) using a Nikon SMZ18. Differences in volume were approximated by differences in area in a single focal plane, with error bars indicating standard deviation.

### Image analysis

Unless otherwise noted, quantitative measures were carried out using FIJI ^63^. Membrane length, cell height and maximal cell width were measured as line length; maximal cell width was measured at the apicobasal position that corresponded to the maximal area in the transverse cell section; columnarization of cells was calculated as the ratio of cell-height and maximal-cell-width. The tapering index (TI) was calculated by subtracting the actual cell volume from the ideal volume of a cell column and then normalizing it to the ideal column volume. The tapering index was determined separately for the apical (from absolute cell apex to 5 µm height) and basal side of the cell (from 5 µm height to absolute basal). The distribution of actin at individual membranes was measured manually along the emerging membrane using the “segmented line” tool in single plane cross-section. Intensities were subsequently plotted using the “plot profile” function. Kymograph heatmaps were generated from a single recording representative of at least three embryos, plotted are mean of five membranes selected at each time point. Plots were generated in Matlab (Version 2016b). The distributions of GFP-Slam and Dia-eGFP within a cell were analyzed by manually defining areas of cell outline (F-actin), nucleus (DNA), and cytoplasm (neither F-actin nor DNA) and then reporting the fluorescent signal indicating GFP-Slam or Dia-eGFP relative to the total measured fluorescence in the same channel per cell. The distributions of GFP-Slam and Dia-eGFP at lateral cell-cell contacts was analyzed in fixed material, where progression of cellularization was binned in three stages based on the linear advance of the cell outline visualized by phallacidin. Bin ranges were adjusted for control and *slam*^OE^ embryos, respectively: early, 0-5 µm and 0-7 µm; mid, 5-8 µm and 7-15 µm; late, 8-13 µm and 15-26 µm. Figure panels were assembled and layouted in Adobe Illustrator.

### Gene expression

To define a set of genes that potentially contributes to *Drosophila* cellularization through substantial increase in zygotic transcription levels, we followed an unbiased approach using previously described microarray data ^21^. The published dataset comprises mRNA expression profiles in biological triplicates that sample the stage before cellularization (T0), during the early (T1) and late phase (T2) of cellularization, and at the onset of (T3) and midway through gastrulation (T4). We used an updated annotation file (DrosGenome1.na35.annot.csv) and identified 821 features with an increase in transcript levels, either at the onset of, or during cellularization as indicated by an average signal log ratio (SLR) of >2 in T0/T1 or T0/T2 comparisons. This list of candidates was narrowed down to 67 features with a dedicated peak during cellularization, which was defined by a subsequent decrease of transcript levels at the onset of gastrulation and an average SLR of <-2 in T1/T3 or T2/T3 comparisons (mean of three replicate comparisons for any transition). For heatmap and cluster analyses, RNA expression levels were compared to the mean expression of all five time points in log2 ratios. Mapping and clustering was performed in Python 3.8 using *seaborn.clustermap* with colormap *coolwarm*, standard parameters, and log2 expression ratios capped at values of <-2 and >2. For each candidate gene, expression in *Chironomus* was tested by a blast search against a transcriptome that comprised all stages of blastoderm formation; if no significant hit could be found, the gene was considered to be not expressed during blastoderm formation in *Chironomus*.

### Phylogenetics and genome analysis

To map gene occurrences onto the dipteran phylogeny, assembled genomes of 171 fly species were retrieved as bulk download from NCBI (tax-id 7147, latest re-run on 09 Sept. 2020; accession numbers listed in **Figure 1 supplement 4**). Individual files were concatenated into one fasta file per species. Quality of genomes was assessed using BUSCO v4.1.2 with dataset odb10_insecta ^64^. BLAST databases were generated using makeblastdb (ncbi-blast-2.10.1+) and searched for putative orthologs using tblastn (word_size 6, gapopen 11, gapextend 1, matrix BLOSUM62). Best hits were used for a reciprocal blast against *Drosophila melanogaster* amnio acid sequences (dmel-all-translation-r6.32.fasta ^65^). E-value cutoff in all blast searches was set to the standard of 0.01 ^66^. Blast searches for *slam* homologues were carried out using NP_001285668. Blast search detection probability for *slam* within the insect order Diptera was carried out using the abSENSE null model of homolog detectability essentially as described ^29^. Briefly, phylogenetic distances within Diptera were computed based on all 171 available fly genomes, which were searched for a set of highly conserved genes using BUSCO v4.1.2 with dataset insecta_obd10 ^64^. As a representation of major dipteran families, 48 species were selected with a set of 25 BUSCO genes that were each present in at least 44 of these genomes. For each gene, a multiple protein sequence alignment was generated using alignment software MUSCLE (version 3.8.31) ^67^ with default parameters. Alignments were then concatenated, the concatenated alignment pruned by trimAl ^68^ with option - *automated1*, and Protdist from the PHYLIP software package (version 3.696) ^69^ was used with default parameters to compute pairwise evolutionary distances for all 48 fly species, measured in substitutions per site. Bitscores for tblastn searches of *slam* were obtained for a representative set of schizophoran species. To account for the difference in search space (nucleotide rather than amino acid databases), database sizes were estimated to 2N, with N being the length of a genome containing ∼N/3 triplets that can be read in six possible reading frames (C Weisman, personal communication). Distances were formatted relative to *Drosophila melanogaster* as focal species and, together with the bitscores, used as input into abSENSE as described ^29^. The graphical detectability prediction result was color adjusted using Adobe Illustrator.

### Statistics

Boxplots were generated by using Excel and statistical comparison were performed via Excel’s implemented students t-test (two sided, unpaired). All samples fit normal distribution; P-values of student’s t-test are indicated in the figure legend. The size of n (embryos and cells) is indicated on each figure.

---

**Figure 1 supplement 1.**
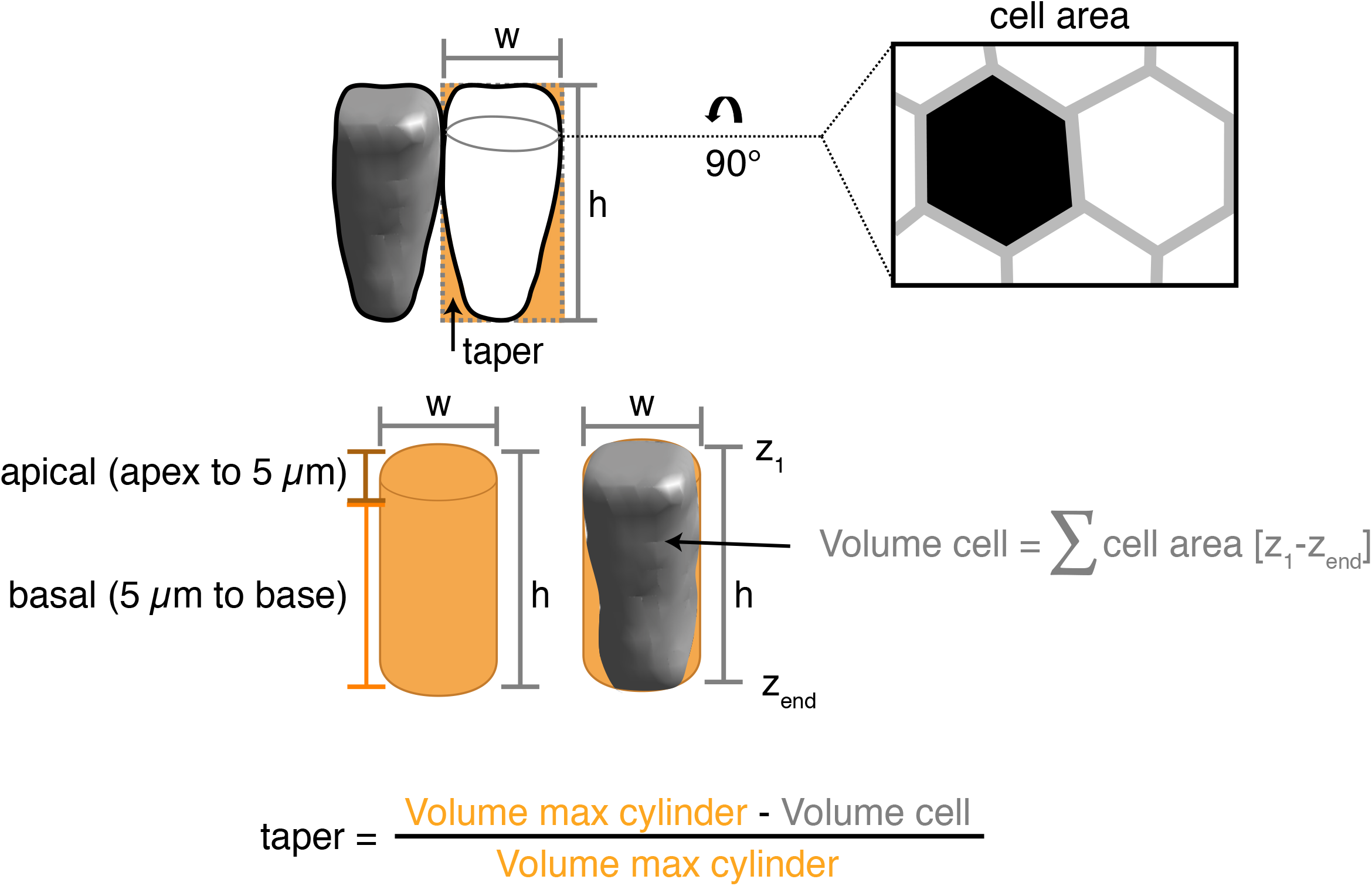
Sketch to illustrate how cell tapering is defined as the deviation of cell shape from an ideal cylinder. The amount of taper was defined as the deviation of the cell volume from the ideal maximal cell column, normalized by the volume of the ideal maximal cell column. Tapering was determined separately for the apical (from absolute cell apex to 5 μm height) and basal side of the cell (from 5 μm height to absolute basal). Cell volume is calculated as the sum of each cell area in a z-stack multiplied with its respective z-stack height.

**Figure 1 supplement 2.**
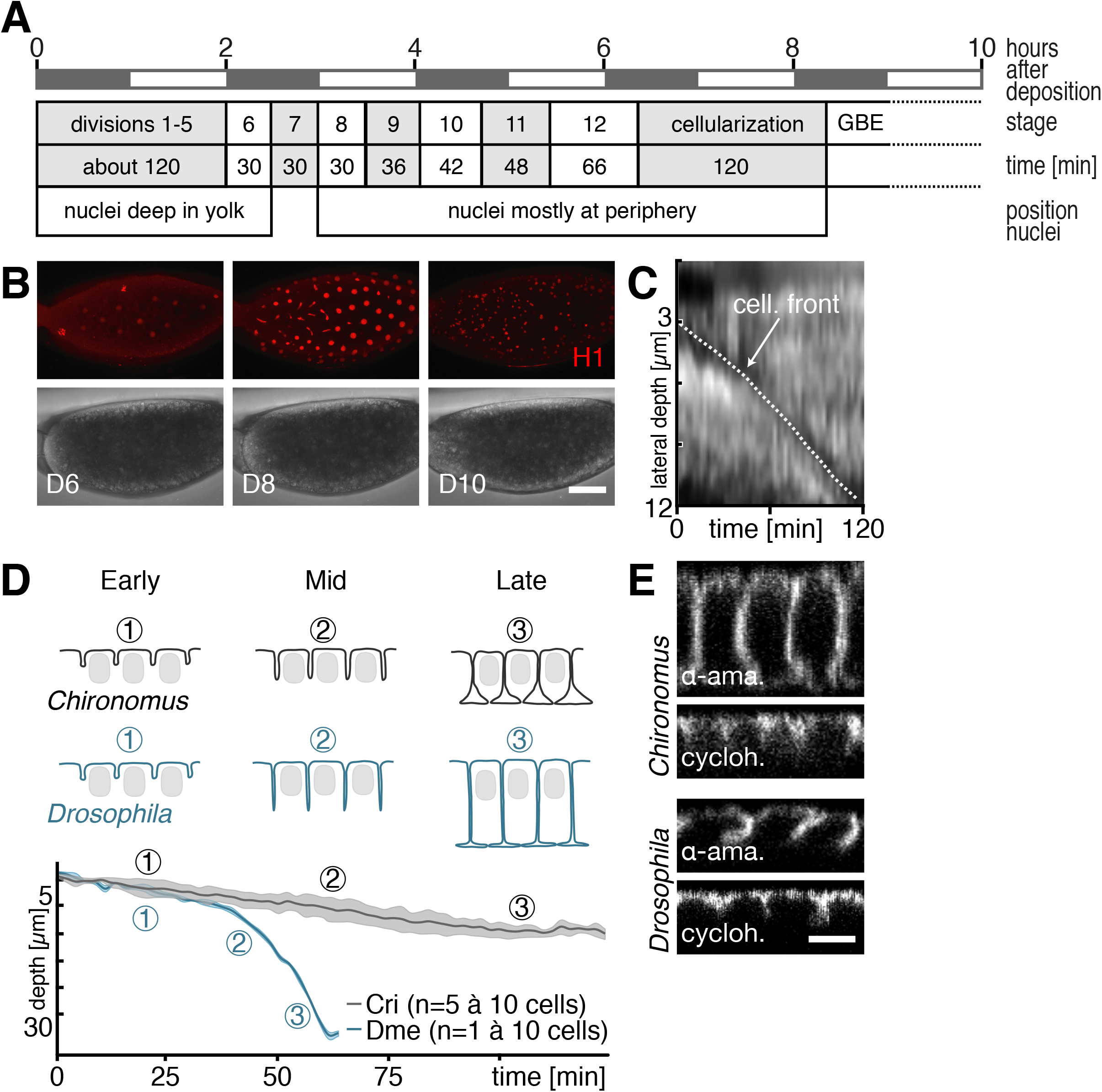
Timing of the first stages of *Chironomus* embryonic development. **A,B**, Timing and duration of nuclear cleavage cycles (A), observed by brightfield and confocal time lapse recordings in embryos injected with fluorescently labeled Histone H1 (B). **C**, Progression of membrane invagination during the course of cellularization in a kymograph derived from DIC recording. **D**, Sketches and measured progression of blastoderm formation in *Chironomus*; the dynamics from a *Drosophila* embryo illustrate previously described dynamics ^19^. **E**, Blastoderm formation in *Chironomus* and *Drosophila* following inhibition of transcription with α-amanitin (α-ama.) and translation with cycloheximide (cycloh.).

**Figure 1 supplement 3.**
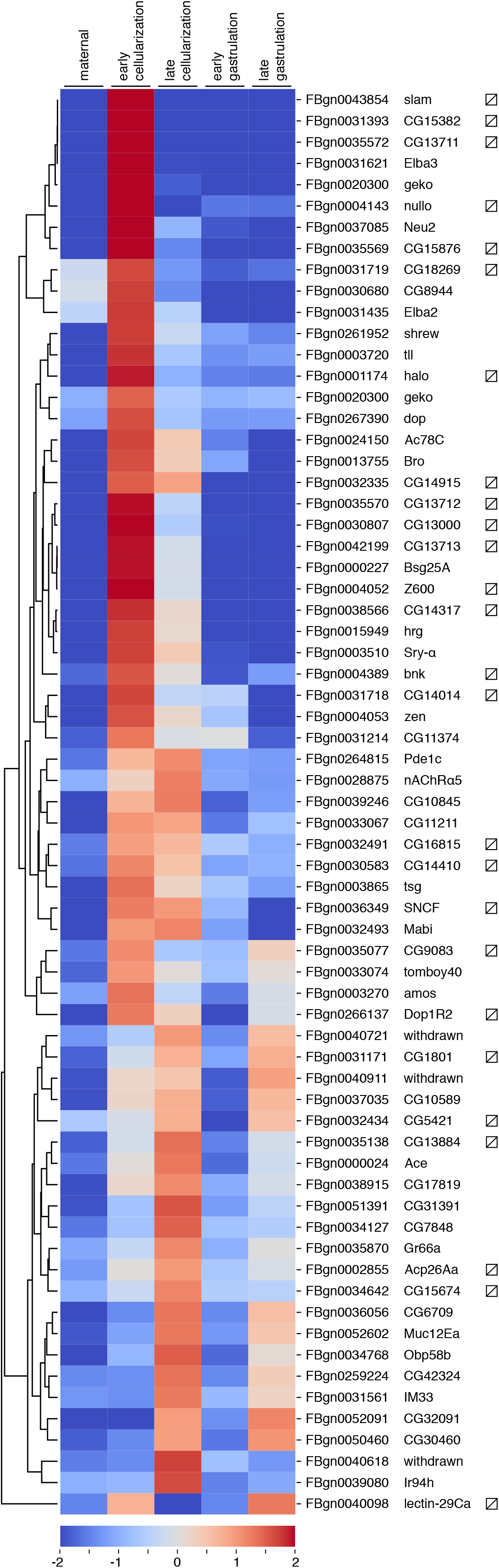
Heatmap cluster representation of 67 features that were selected from a described Affymetrix microarray dataset ^21^ based on significant upregulation during *Drosophila* blastoderm formation (left). Indicated are corresponding flybase identifier and gene names based on current annotation file (middle). Candidates for further analyses identified by absence of significant blast hit in zygotic *Chironomus* transcriptome (strike-through box, right). Stages (maternal, early cellularization, late cellularization, early gastrulation, late gastrulation) correspond to stages T0-T4 as described ^21^. Shown for each feature are increased (red) and decreased (blue) RNA expression levels compared to the mean expression of all five time points in log2 ratios.

**Figure 1 supplement 4.**
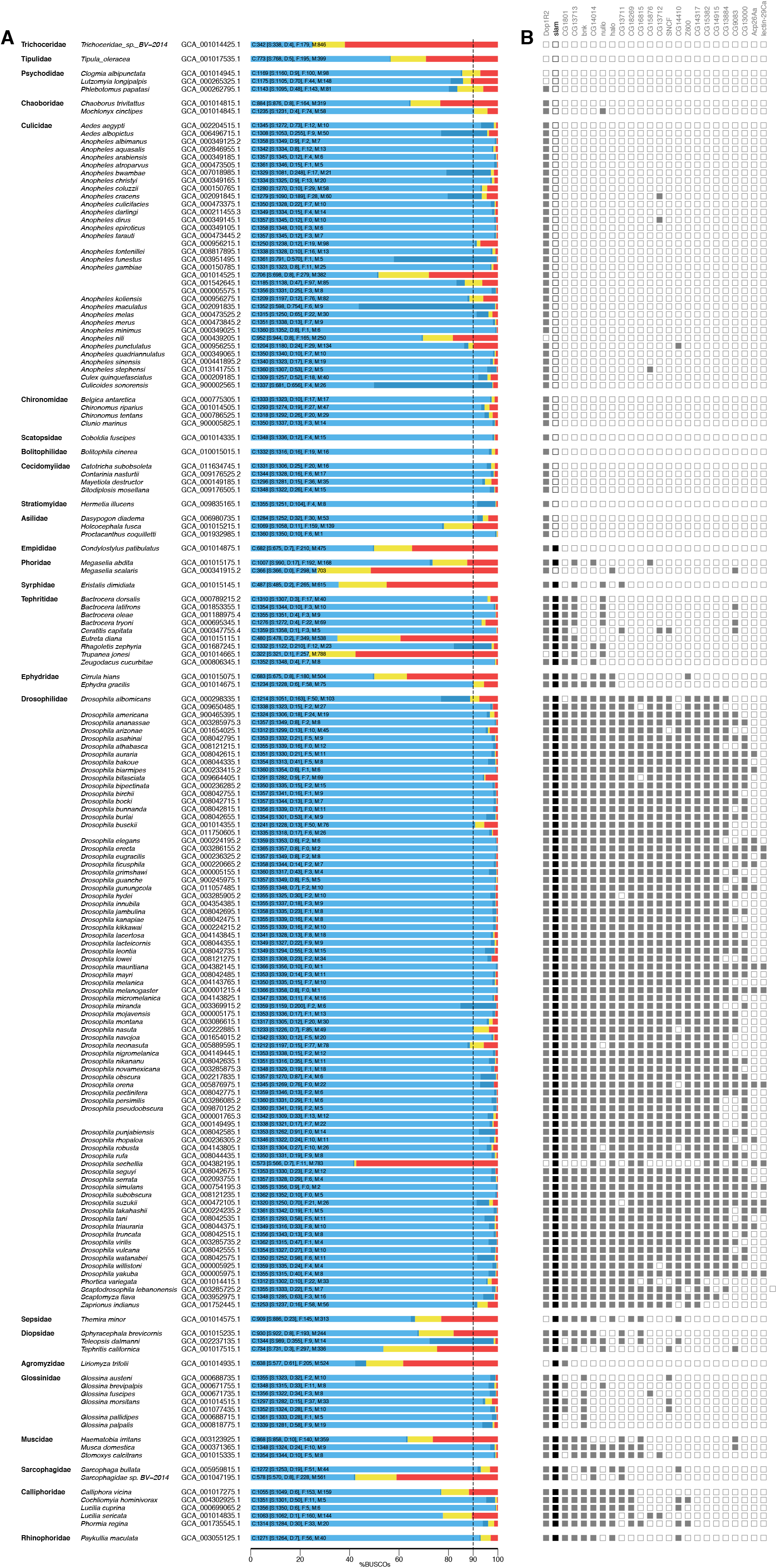
Analysis of fly genomes integrity and search for orthologs. **A**, List is grouped by fly families and indicates species name followed by NCBI accession number of each genome analyzed. Indicated are proportions of complete and single copy genes (light blue), of complete and duplicated genes (dark blue), of fragmented genes (yellow), and missing genes (red) in BUSCO set insecta_odb10. Indicated is whether a *slam* ortholog could (filled box) or could not (empty box) be identified by reciprocal blast searches. Genomes with less than 10% of missing BUSCO genes were considered as “high integrity” genomes and counted accordingly in Figure 1. **B**, Identification of orthologs for candidate genes (top) by reciprocal blast searches indicated by filled box.

**Figure 1 supplement 5.**
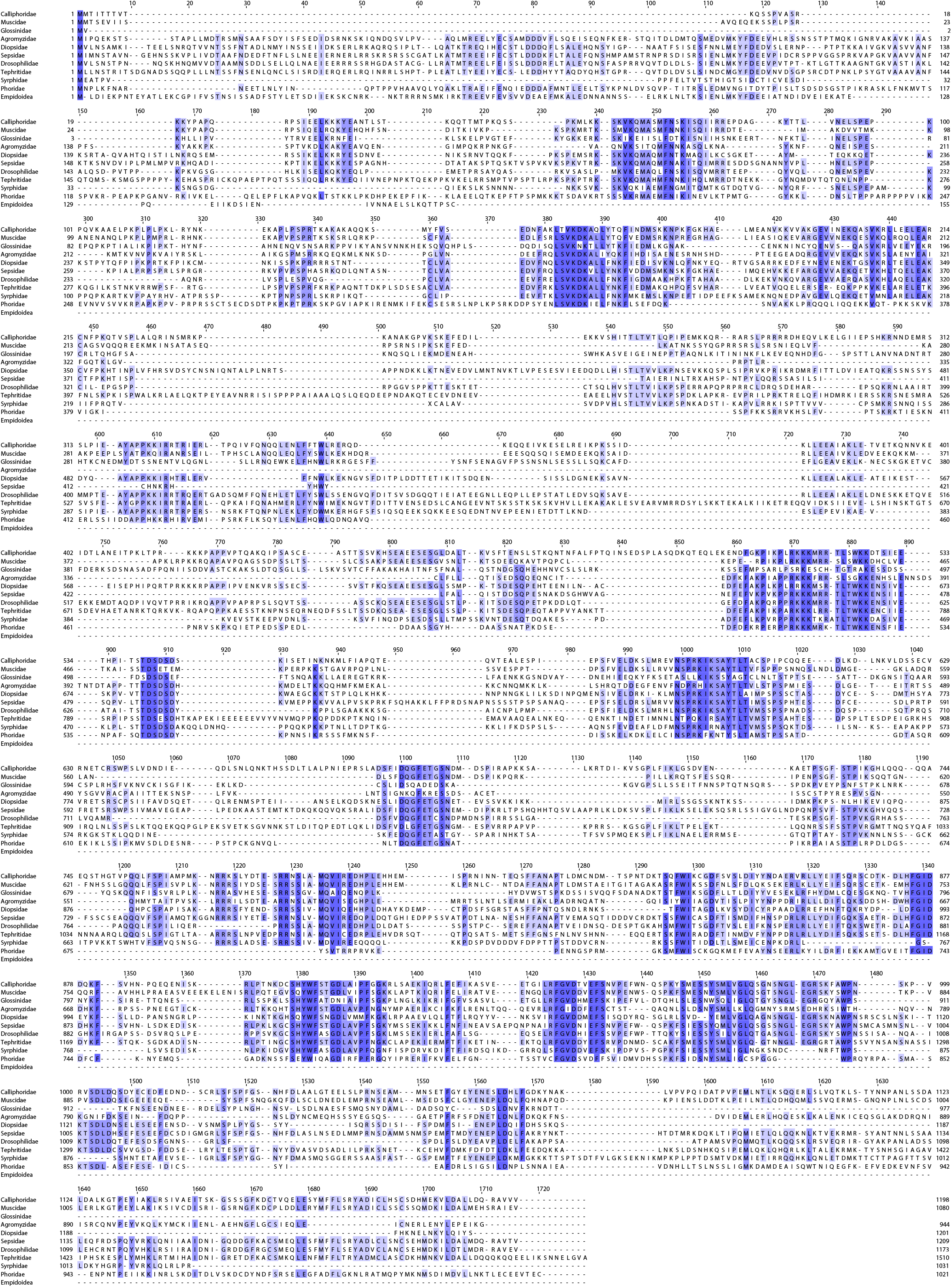
Protein alignment of putative Slam sequences in species representing the major cyclorrhaphan fly families. Sequences are from *Lucilia cuprina* (Calliphoridae), *Musca domestica* (Muscidae), *Glossina pallidipes* (Glossinidae), *Liriomyza trifolii* (Agromyzidae), *Teleopsis dalmanni* (Diopsidae), *Themira minor* (Sepsidae), *Drosophila melanogaster* (Drosophilidae), *Bactrocera dorsalis* (Tephritidae), *Episyrphus balteatus* (Syrphidae), *Megaselia abdita* (Phoridae), and *Condylostylus patibulatus* (Empidoidea).

**Figure 1 supplement 6.**
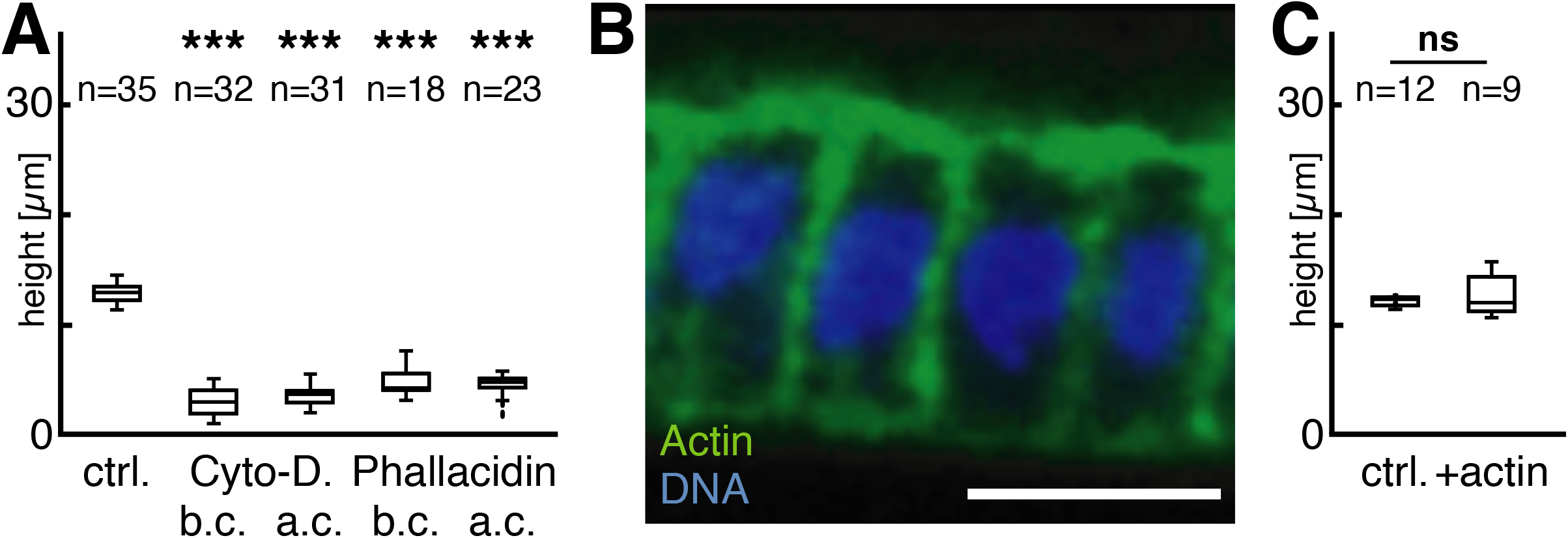
**A**, Impaired blastoderm formation after injection of toxins affecting F-actin stability (Cyto-D, Cytochalsin D; Phal, Phallacidin) before (b.c.) and after (a.c.) onset of cellularization. Statistical significance computed in comparison to control injected embryos (ctrl; \*\*\**P*<0.0001). **B,C**, Cell height (B) not affected by injection of mouse actin monomers (C; ns, *P*=0.201). Scale bars 10 μm.

**Figure 2 supplement 1.**
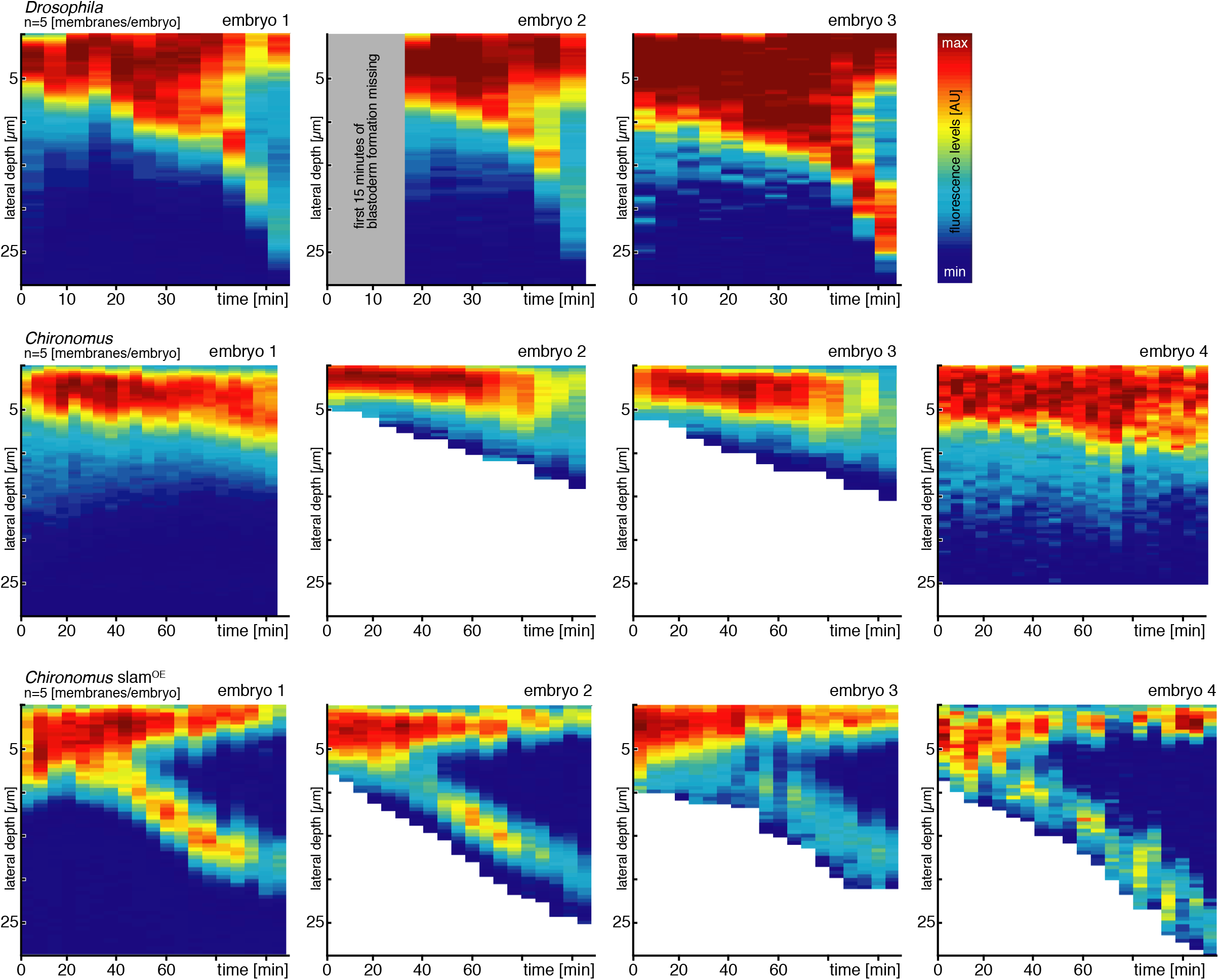
Heatmap kymographs of additional recordings as in main Figure 2B,D,J, illustrating local enrichment of F-actin during blastoderm formation in *Drosophila* wildtype (top)*, Chironomus* wildtype (middle), and *Chironomus slam*^OE^ embryos (bottom). For reference, the first embryo recordings correspond to the kymographs in Figure 2.

**Figure 3 supplement 1.**
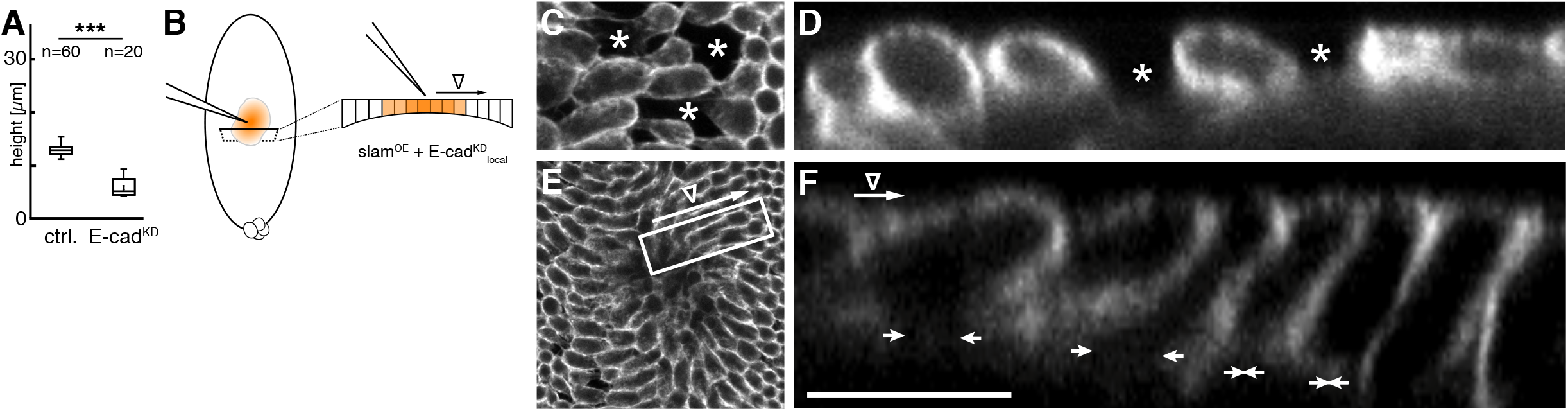
Effect of *E-cad* knockdown on *Chironomus* blastoderm formation. **A**, In embryos injected with dsRNA against *Cri-E-cad* during nuclear migration stage, blastoderm formation stalls (*** *P*<0.0001). **B**, Sketch outlining the experimental approach to study the effect of local *Cri-E-cad* knockdown in *slam*^OE^ embryos. **C,D**, Blastoderm in a representative *Cri-E-cad*^KD^_local_ *slam*^OE^ embryo at site of maximal *E-cad*^KD^ effect, with typical signs of cell individualization (asterisks) within the blastoderm in areas of poor adhesion, shown as a maximum intensity projection en face (C) and in a xz-reslice (D). **E-F**, Graded response to local *Cri-E-cad* knockdown in *slam*^OE^ embryo shown en face (E) and in a xz-reslice of a selected window (F). The inferred activity gradient (∇) of *Cri-E-cad* knockdown is indicated with long arrow, short arrows indicate completion of blastoderm formation as cell base is open or closed. Scale bar is 10 μm.

**Figure 3 supplement 2.**
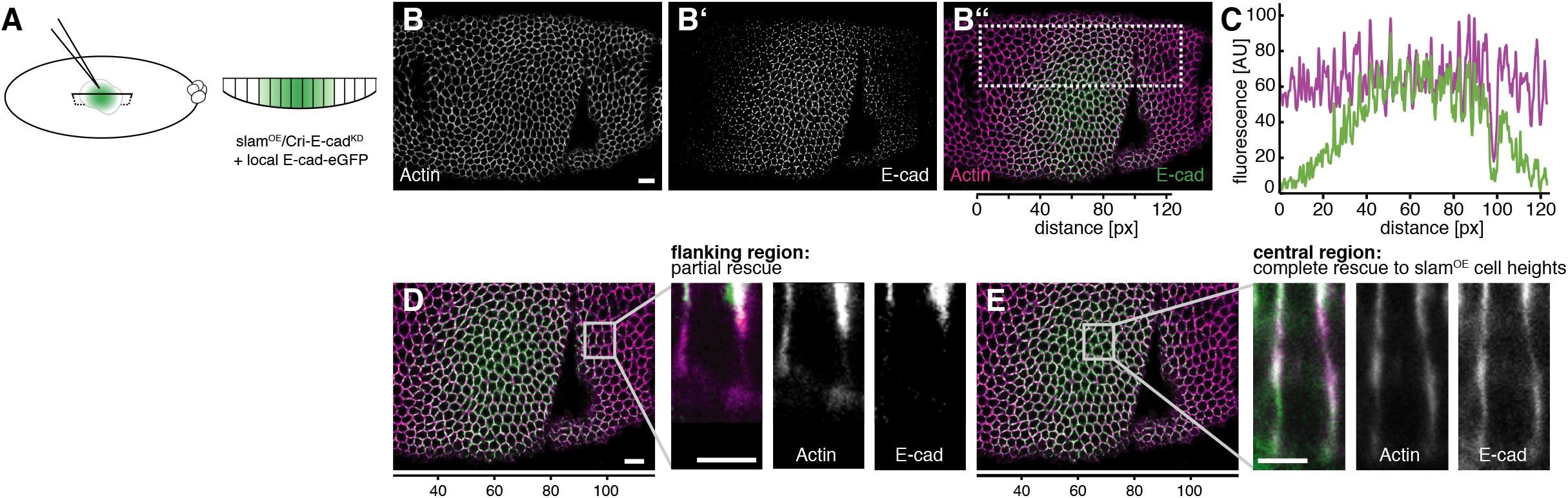
Local E-cad rescue of blastoderm formation in *slam*^OE^/*Cri-E-cad* RNAi embryos. **A**, Sketch outlining the experimental approach to study the effect of local E-cad availability in *slam*^OE^/*Cri-E-cad* RNAi embryos. **B-B”**, F-actin and E-cad levels visualized by fluorescent reporters (Lifeactin-mCherry and E-cad-eGFP). **C**, Quantification of fluorescent signal in subapical plane (3-5 μm) over the area shown in B’’. **D**, Distribution of F-actin and E-cad in a cell (shown in a xz-reslice) representative of the blastoderm flanking maximal E-cad levels seen in (C). **E**, Distribution of F-actin and E-cad in a cell representative of the blastoderm with maximal E-cad levels. Scale bar is 5 μm.

## Notes

### Competing Interest Statement

The authors have declared no competing interest.

